# IL-10 Overexpression Improves Cerebral Microcirculation and Attenuates Cerebral Vasospasm After Experimental SAH

**DOI:** 10.64898/2026.08.25.747167

**Authors:** Kenshu Nogami, Hiroshi Ishii, Munehiro Demura, Takuya Nakamura, Nguyen Dinh Loc, Mika Takarada-Iemata, Yuji Tsunekawa, Yuko Nitahara-Kasahara, Takashi Okada, Tomoya Kamide, Mitsutoshi Nakada, Osamu Hori

## Abstract

**BACKGROUND:** Subarachnoid hemorrhage (SAH) induces inflammatory responses and subsequent immune cell activation, which may contribute in cerebral vasospasm, microcirculatory impairment and poor neurological outcomes. Although cerebral vasospasm has traditionally been considered a major cause of delayed cerebral ischemia after SAH, therapies targeting angiographic vasospasm have not consistently improved functional outcomes. Early inflammatory responses may contribute to microcirculatory impairment, cerebral vasospasm, and subsequent neurological injury. Herein, we investigated whether interleukin-10 (IL-10), an anti-inflammatory cytokine, improves these outcomes in an experimental SAH model.

**METHODS:** Mice received intramuscular injections of either an adeno-associated virus encoding IL-10 (AAV/IL-10) vector or an AAV expressing green fluorescent protein (AAV/GFP) vector (control). India ink angiography was performed to assess the diameter of the sphenoidal segment of the middle cerebral artery (MCA), the total length of the visible cortical arteries, and cortical staining intensity, as indices of cerebral vasospasm, microcirculatory impairment, and cerebral perfusion, respectively. Perivascular inflammatory cell infiltration and cytokine levels were assessed using immunohistochemistry and ELISA. We also evaluated the therapeutic efficacy of the AAV/IL-10 vector when administered immediately after SAH induction.

**RESULTS:** IL-10 overexpression significantly improved neurological outcomes after SAH and was associated with attenuated cerebral vasospasm and microcirculatory impairment, as well as preservation of cerebral perfusion. It also significantly reduced neutrophil and macrophage infiltration around the internal carotid artery and attenuated SAH-induced elevations in IL-6 and matrix metalloproteinase-3 levels. Mice treated with the AAV/IL-10 vector immediately after SAH induction showed significant improvements in neurological scores and cerebral perfusion.

**CONCLUSIONS:** AAV-mediated IL-10 overexpression improves neurological outcomes after SAH, likely by attenuating inflammatory responses, cerebral vasospasm, and microcirculatory impairment. These findings suggest that IL-10-based anti-inflammatory therapy is a promising therapeutic strategy for SAH.

## Introduction

Despite advances in surgical and neurocritical care management, aneurysmal subarachnoid hemorrhage (SAH) remains a devastating form of stroke associated with high mortality and long-term neurological disability.^1,2^ Cerebral vasospasm of the major arteries, such as the internal carotid artery (ICA) and middle cerebral artery (MCA), has long been considered a major cause of delayed cerebral ischemia (DCI) and poor outcomes after SAH.^3^ However, therapeutic strategies that primarily target angiographic vasospasm have not consistently improved functional outcomes, suggesting that earlier and more complex pathological processes contribute to brain injury after SAH.^4^

Accordingly, increasing attention has focused on the sequential pathological cascade after SAH, in which early brain injury (EBI) is driven by microcirculatory impairment, subsequently leading to DCI and ultimately determining long-term neurological outcomes. Microvascular constriction, microthrombosis, endothelial dysfunction, and inflammatory cell activation occur shortly after hemorrhage during the EBI phase.^5–7^ We recently reported that neutrophil- mediated receptor for advanced glycation end product (RAGE) signaling contributes to cortical microcirculatory impairment and cerebral vasospasm after experimental SAH.^8^ Combined with clinical evidence that reduced soluble RAGE levels predict symptomatic vasospasm,^9^ these findings suggest that early inflammatory responses may drive a pathological sequence from microcirculatory impairment to cerebral vasospasm and DCI.

Recent studies have demonstrated that neutrophils and macrophages play central roles in the pathogenesis of brain injury after SAH.10 Blood components and damage-associated molecular patterns entering the subarachnoid space activate immune cells such as neutrophils and macrophages, thereby promoting the production of pro-inflammatory cytokines and amplifying the inflammatory response.5,9,11 This inflammatory cascade disrupts the structural integrity of the blood–brain barrier (BBB) and increases vascular permeability, leading to increased neutrophil and macrophage infiltration into the affected area.12 Activated neutrophils further amplify the inflammatory response through the release of proteolytic enzymes, such as myeloperoxidase (MPO), and the formation of neutrophil extracellular traps (NETs), which contribute to endothelial injury, microthrombus formation, cerebral microcirculatory impairment, and cerebral vasospasm.13-15 Ultimately, these processes result in neuronal injury and neurological dysfunction.

Given the pivotal role of inflammation in the development of cerebral vasospasm and microcirculatory impairment after SAH, modulation of inflammatory responses represents a promising therapeutic strategy. Interleukin-10 (IL-10) is a key endogenous anti-inflammatory cytokine that regulates immune resolution by suppressing the production of pro-inflammatory mediators such as interferon-γ, tumor necrosis factor-α, IL-1β, and IL-6.^16,17^ Therapeutic effects of IL-10 administration have been demonstrated in experimental models of cerebral ischemia.^18–22^ Furthermore, Nakajima et al. demonstrated that sustained systemic overexpression of IL-10 using an adeno-associated virus (AAV) vector improved neurological dysfunction in ischemic stroke models.^23^

Therefore, we tested the hypothesis that AAV-mediated IL-10 overexpression improves cerebral vasospasm, microcirculatory impairment, and neurological outcomes by regulating inflammatory responses in a mouse model of SAH.

## METHODS

### Data Availability

Data will be made available upon reasonable request. Please refer to the Major Resources Table in the Supplemental Materials.

### Animals

All animal experiments were approved by the Animal Care and Use Committee of Kanazawa University (approval number: AP-224352) in accordance with the Fundamental Guidelines for Proper Conduct of Animal Experiments and Related Activities in Academic Research Institutions under the jurisdiction of the Ministry of Education, Culture, Sports, Science and Technology (Japan) and in compliance with the ARRIVE 2.0 guidelines. A total of 127 C57BL/6 male mice were used in this study. The mice were purchased from Japan SLC (Hamamatsu, Shizuoka, Japan) and housed in standard cages under a 12 h light/dark cycle with food and water available ad libitum.

### Study Design, Randomization, and Blinding

Animals were randomly assigned to their respective experimental groups before intervention or SAH induction. To minimize bias, all outcome assessments, including neurobehavioral testing, image analysis for India ink angiography, histological cell counting, and biochemical analyses (ELISA and cytokine arrays), were performed by two or more investigators (K.N., T.N., and H.I.) who were blinded to group allocation.

Sample sizes were predetermined based on preliminary and pilot data. Power analyses performed using G*Power v3.1.9.6 indicated that a sample size of five animals per group provided >92% statistical power (92.1%–98.1%) to detect significant differences across all primary neurobehavioral, histological, and biochemical endpoints at an alpha level of 0.05.

### AAV Vector Production, Purification, and Quantification

Myo-AAV/IL-10 and Myo-AAV expressing green fluorescent protein (AAV/GFP) vectors were generated using an AAV-MAX Transfection Kit (Thermo Fisher Scientific) in VPC2.0 cells. Following cell lysis, the vectors were purified by affinity chromatography using an ÄKTA avant 25 system (Cytiva). Purified AAV vector genome titers were quantified by real- time PCR using the AAVpro® Titration Kit Ver.2 (Takara Bio). AAV vectors were purified as previously described.^24^ Detailed procedures for plasmid transfection, purification, and quantification are provided in the Supplemental Materials.

### Administration of AAV Vectors *in Vivo*

Mice received intramuscular injections of AAV vectors into the left biceps femoris muscle. Mice in the control group received the Myo-AAV/GFP (AAV/GFP) vector (1.0 × 10¹² genome copies [gc] in 30 μL), whereas mice in the IL-10 overexpression group received the Myo-AAV encoding IL-10 (AAV/IL-10) vector (1.0 × 10¹² gc in 5 μL). Although the injection volumes differed between groups, the total viral dose was standardized to 1.0 × 10¹² gc in both groups.

### Blood Collection and Measurement of Serum IL-10 Levels in Mice

To evaluate the time course of serum IL-10 levels, 7-week-old mice received the AAV/IL- 10 vector. Blood samples (300 μL) were collected from the submandibular vein under isoflurane anesthesia. For serum preparation, blood samples were allowed to stand at room temperature for 1 h after collection, centrifuged at 2,000 × g for 20 min at 20 °C, and the supernatant was transferred to fresh tubes and stored at −80 °C until analysis. Serum IL-10 concentrations were measured using a mouse IL-10 ELISA kit (R&D Systems), according to the manufacturer’s instructions. Twenty mice were used in this study.

### Animal Model of SAH

To investigate the effects of AAV-mediated IL-10 overexpression on SAH, SAH was induced 5 weeks after AAV vector administration in 7-week-old mice (Figure 1A). To evaluate the therapeutic efficacy of AAV vector administration after SAH, SAH was induced in 12- week-old mice and immediately followed by AAV vector administration (Figure 5A). To avoid the potential influence of blood collection on outcomes after SAH, blood was not collected from mice subjected to SAH induction following AAV vector administration. The SAH model was established using the endovascular perforation method as previously described (Figure 1B and Supplemental Methods).^25^ As surgical controls, AAV/GFP sham and AAV/IL-10 sham mice were included. These mice underwent all surgical procedures except arterial perforation, and all were included in the final analyses. SAH severity was assessed using the grading system described by Sugawara et al.^26^

**Figure 1.**
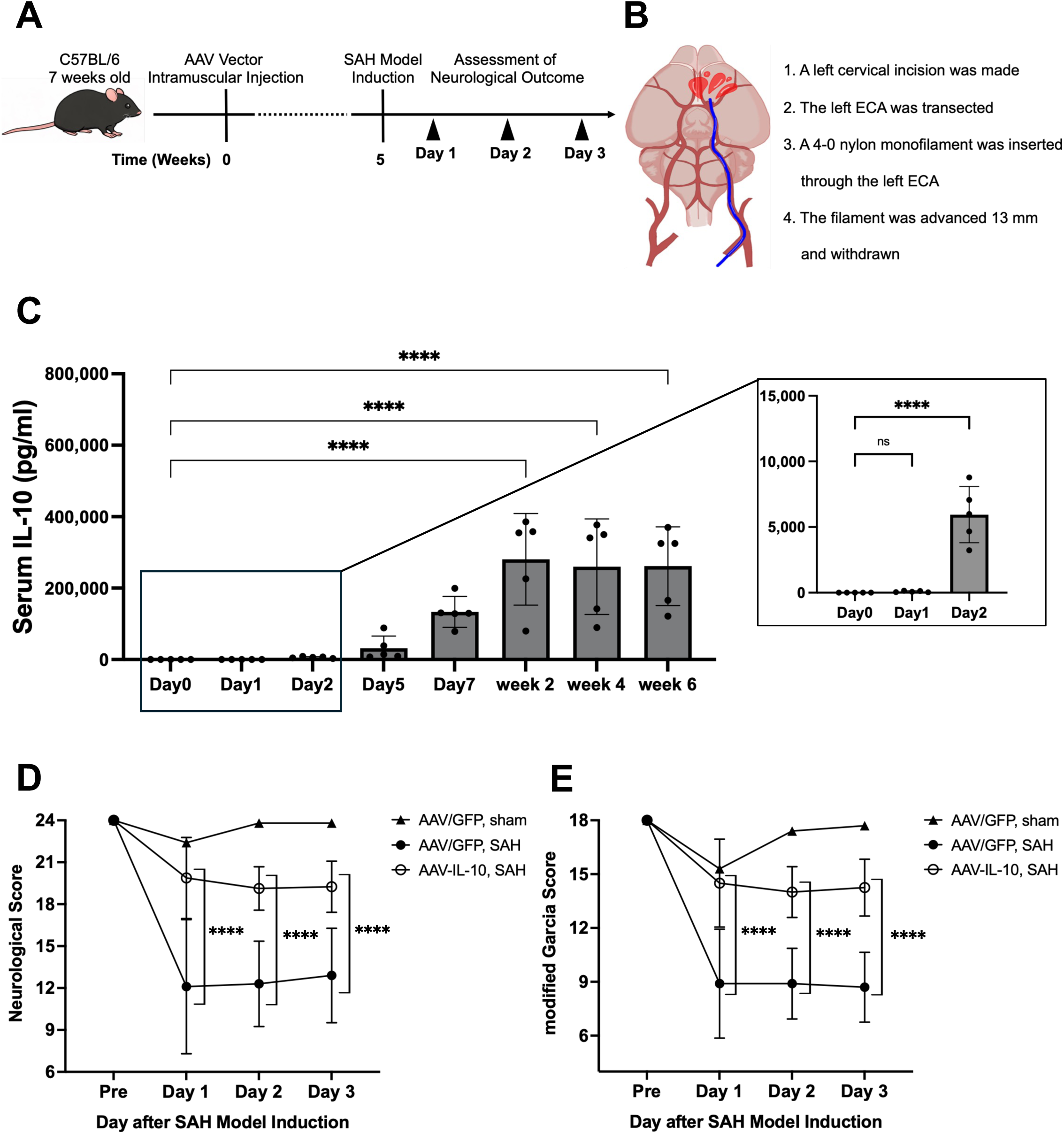
Interleukin-10 (IL-10) Overexpression Improves Neurological Outcomes After Experimental Subarachnoid Hemorrhage (SAH) A, Experimental protocol. The SAH model was established 5 weeks after adeno-associated virus (AAV) vector administration. B, Endovascular perforation method. C, Time course of serum IL-10 concentrations after AAV/IL-10 administration. Serum IL-10 levels reached a plateau 2 weeks after vector administration. Values are presented as mean ± SD, n = 5. D and E, Neurological outcomes after SAH assessed using the neurological and modified Garcia scores. Values are presented as mean ± SD, n = 10 in the AAV expressing green fluorescent protein (AAV/GFP) sham, AAV/GFP SAH, and AAV/IL-10 SAH groups. **** *P* < 0.0001 versus the AAV/GFP SAH group.

### Neurobehavioral Testing

Neurological outcomes were assessed using two tests at baseline (before SAH) and on days 1–3 after SAH (Figure 1A and 5A). Six mice (two AAV/GFP SAH, three AAV/IL-10 SAH, and one AAV/IL-10 post-SAH) that died within 3 days after SAH were excluded from analysis. The modified Garcia score (range, 3–18 points) is a composite score consisting of six items: spontaneous activity, spontaneous movement of all limbs, forelimb movement, climbing, side- stroking response, and vibrissa touch response. Higher scores indicate better neurological outcomes.^27^ The neurological score (range, 8–24 points), which evaluates sensorimotor function, was calculated as the sum of eight items: spontaneous activity, climbing, balance, side-stroking response, vibrissa touch response, visual response, forelimb use, and hindlimb use. Similarly, higher scores indicate better neurological outcomes.^28^ A total of 35 mice were used for the experiments shown in Figure 1 and six mice for those shown in Figure 5.

### India Ink Angiography

Intracranial vessels were assessed by India ink angiography 48 h after SAH or sham surgery, as previously reported.^29^ Perfusion was performed under a controlled pressure of 60– 80 mmHg, which approximates the physiological blood pressure of mice.^30^ Detailed procedures are described in the Supplemental Methods. After perfusion, animals were maintained at 4 °C for 24 h. The brains were then harvested, and the circle of Willis, basilar artery, and cortical vessels were photographed at constant magnification using a stereomicroscope (SMZ800-1, Nikon) equipped with a digital camera (COOLPIX4500, Nikon). The images were analyzed using ImageJ software (National Institutes of Health) (Figure S1A). Microcirculatory impairment was assessed by quantifying the total length of India ink-filled cortical arteries, whereas cerebral vasospasm was evaluated by measuring the diameter of the left MCA (Lt MCA), and cerebral perfusion was assessed by determining the cerebral perfusion index (CPI). Detailed evaluation methods are described in the Supplemental Methods. Forty and six mice were used in the experiments shown in Figures 2 and 5, respectively. Seven mice (three AAV/GFP SAH, three AAV/IL-10 SAH, and one AAV/IL-10 post-SAH) that died within 48 h after SAH were excluded from analysis.

**Figure 2.**
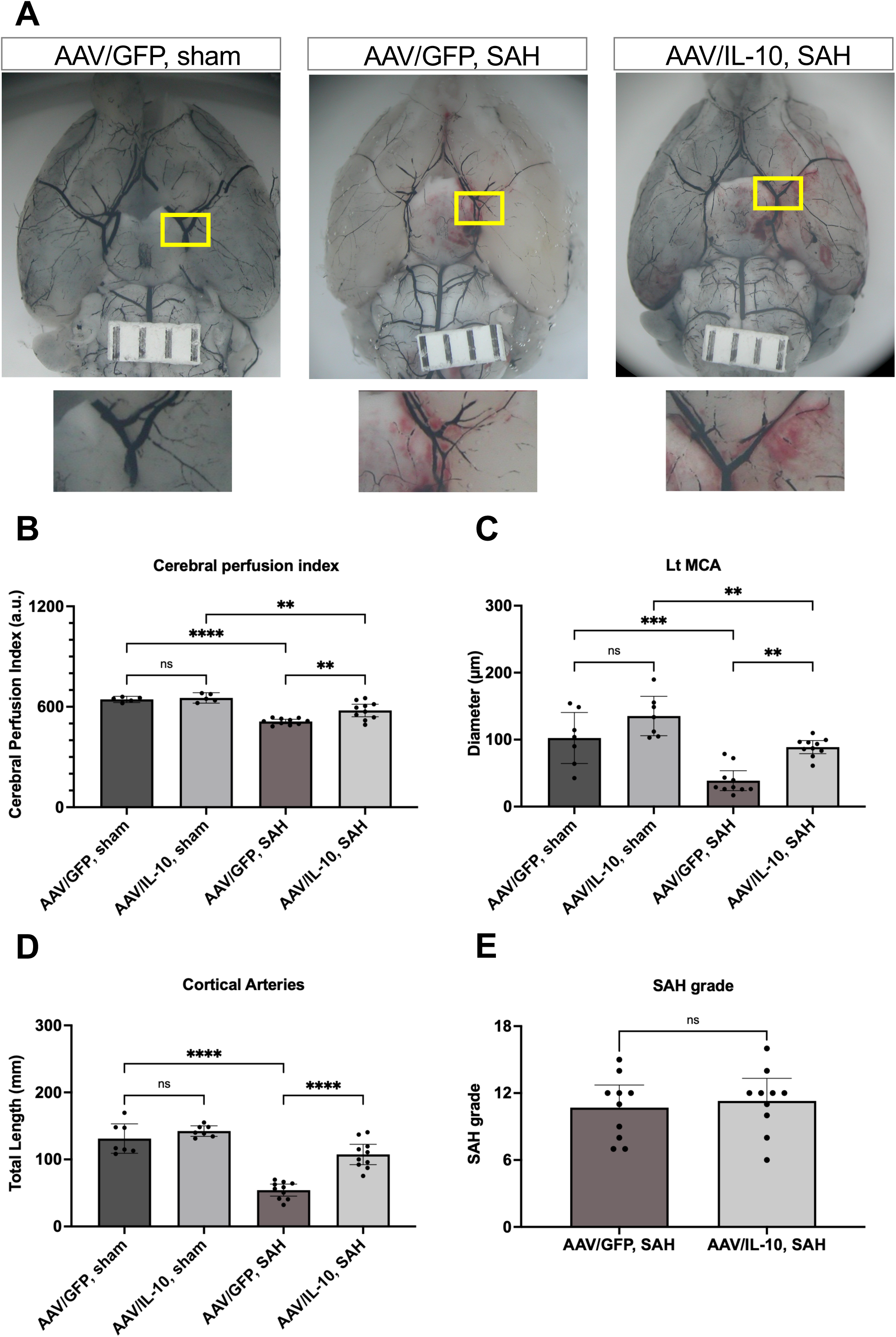
Interleukin-10 (IL-10) Improves Cerebral Microcirculation and Attenuates Cerebral Vasospasm After Experimental Subarachnoid Hemorrhage (SAH). A, Representative India ink angiograms obtained 48 h after SAH in the adeno-associated virus expressing green fluorescent protein (AAV/GFP) sham, AAV/GFP SAH, and AAV/IL-10 SAH groups. Compared with the AAV/GFP SAH group, the AAV/IL-10 SAH group showed enhanced India ink perfusion in the cerebral cortex, indicating improved cerebral microcirculation. Scale bar, 1 mm. B–D, Quantification of India ink staining. E, SAH grade. IL-10 significantly increased the cerebral perfusion index (B), left middle cerebral artery (Lt MCA) diameter (C), and total cortical arterial length (D). Values are presented as mean ± SD, n = 7 in the AAV/GFP sham and AAV/IL-10 sham groups and n = 10 in the AAV/GFP SAH and AAV/IL-10 SAH groups. \*\**P* < 0.01, \*\*\**P* < 0.001, \*\*\*\**P* < 0.0001.

### Immunohistochemistry and Quantification of Perivascular Neutrophils and Macrophages around the Ruptured Vessel

At 48 h after SAH induction, mice were deeply anesthetized and transcardially perfused with phosphate-buffered saline followed by 4% paraformaldehyde, after which the brains were carefully harvested. The specimens were additionally fixed in 4% paraformaldehyde for 2 h and subsequently cryoprotected in 30% sucrose at 4 °C for a minimum of 24 h. Cortical 10- μm-thick cryosections were then prepared. Immunofluorescence staining was performed using primary antibodies against MPO (R&D Systems) and ionized calcium-binding adaptor molecule 1 (Iba1; FUJIFILM Wako). Fluorescent labeling was visualized using Alexa Fluor 488 (cat. A11055) or Cy3-conjugated secondary antibodies (Jackson ImmunoResearch Laboratories, cat. 111-165-003). Staining procedures followed previously established protocols.^7^ Representative images were obtained using a laser scanning confocal microscope (Eclipse TE2000U; Nikon, Tokyo, Japan) at ×200 magnification with Nikon EZ-C1 software. Images were acquired using a fluorescence microscope (BZ-X1000; KEYENCE, Osaka, Japan). Quantitative assessment of perivascular neutrophil and macrophage infiltration was performed in the subarachnoid space of coronal sections containing the left ICA corresponding to the perforation side. Neutrophils were defined as MPO-positive/Iba1-negative cells, whereas macrophages were defined as Iba1-positive cells.

Within the segment extending 1 mm proximally from the bifurcation of the ICA into the middle and anterior cerebral arteries, five representative sections showing the greatest neutrophil accumulation were selected for analysis. Cell analysis was performed using BZ-X Analyzer software (KEYENCE). The average neutrophil and macrophage counts per section were used for statistical evaluation. Ten mice were used in this experiment. No mice were excluded from the experiment.

### Proteome Cytokine/Chemokine Array

For cytokine array analysis, blood samples were collected from mice subjected to experimental SAH. Blood was obtained by direct cardiac puncture at 3 h after SAH, allowed to stand at room temperature for 1 h, and then centrifuged at 2,000 × g for 20 min at 20 °C. Serum was collected and stored at −80 °C until analysis. Relative serum levels of cytokines and chemokines after SAH were evaluated using the Proteome Profiler™ Array (Mouse Cytokine Array, Panel A; R&D Systems Inc.). The membranes were imaged and analyzed using a LuminoGraph I imaging system (ATTO, Tokyo, Japan) and ImageJ software (National Institutes of Health). To further quantify matrix metalloproteinase-3 (MMP-3), which showed a difference in spot intensity on the array, ELISA was performed using the same samples. Additionally, serum levels of IL-6, a representative inflammatory cytokine, were measured using the same samples. Serum concentrations of MMP-3 and IL-6 were determined using mouse MMP-3 and IL-6 ELISA kits (R&D Systems), respectively, according to the manufacturer’s instructions. Ten mice were used in this experiment. No mice were excluded from the experiment.

### Statistical Analysis

Statistical analyses were performed using GraphPad Prism version 11.0.0 (GraphPad Software). Normality was tested using the Shapiro–Wilk normality test. If the data were normally distributed, two-group comparisons were performed using an unpaired t-test with Welch’s correction. If the data were not normally distributed, the Mann–Whitney U test was used. Multiple-group comparisons were performed using one-way analysis of variance (ANOVA) followed by Tukey’s multiple-comparison test. Time-course analyses of neurological scores were performed using two-way ANOVA followed by Bonferroni’s post hoc test.

Statistical power was calculated using G*Power v3.1.9.6. Data are presented as dot plots with bars indicating the mean ± standard deviation (SD). Two-tailed *P < 0.05, **P < 0.01, and ***P < 0.001 were considered statistically significant.

## RESULTS

### IL-10 Overexpression Attenuated Neurological Deficits After SAH

We measured temporal changes in serum IL-10 levels after administration of the AAV/IL- 10 vector. Following intramuscular injection of the AAV/IL-10 vector, serum IL-10 levels began to increase, exceeding known therapeutic concentrations by day 2,^23,31,32^ and continued to increase until the mean serum IL-10 concentration reached a plateau at 2 weeks (Figure 1C). Elevated IL-10 levels were sustained throughout the observation period, confirming stable systemic overexpression of IL-10 following AAV-mediated gene delivery. In contrast, mice in the AAV/GFP group showed no detectable increase in serum IL-10 levels.

Experimental SAH was induced in AAV/GFP and AAV/IL-10 mice, and neurological outcomes were compared between the two groups using the neurological and modified Garcia scores. Both scores decreased on days 1, 2, and 3 after SAH; however, these reductions were significantly attenuated in the AAV/IL-10 SAH group compared with the AAV/GFP SAH group (Figure 1D and 1E). Thus, IL-10 overexpression attenuated neurological deficits after SAH.

### IL-10 Overexpression Attenuated the Decrease in Cerebral Perfusion After SAH

Because IL-10 overexpression attenuated neurological deficits after SAH, we next examined its effects on cerebral perfusion. India ink angiography performed 48 h after SAH demonstrated greater cortical staining compared with the AAV/GFP SAH group (Figure 2A). Consistently, the CPI was significantly preserved in the AAV/IL-10 SAH group (Figure 2B). Cerebral vasospasm, assessed by the Lt MCA diameter, was significantly attenuated in the AAV/IL-10 SAH group compared with the AAV/GFP SAH group (Figure 2C). Similarly, microcirculatory impairment, evaluated by the total length of India ink-perfused cortical arteries, was significantly attenuated in the AAV/IL-10 SAH group (Figure 2D). In contrast, SAH severity did not differ between the two groups (Figure 2E). These findings suggest that IL-10 overexpression attenuated the decrease in cerebral perfusion after SAH by reducing major arterial vasospasm and microcirculatory impairment.

### IL-10 Reduced Neutrophil and Macrophage Accumulation around the Left ICA After SAH

Perivascular accumulation of neutrophils and macrophages around cerebral arteries in the subarachnoid space after SAH has been reported previously.^33^ To investigate how IL-10 overexpression influences the accumulation of these inflammatory cells, we performed immunohistochemical staining using an anti-MPO antibody as a neutrophil marker and an anti- Iba1 antibody as a macrophage marker. In the AAV/GFP SAH group, accumulation of MPO-positive neutrophils and Iba1-positive macrophages was observed around the left ICA 48 h after SAH (Figure 3A). In contrast, neutrophil and macrophage accumulation was significantly reduced in the AAV/IL-10 SAH group (Figure 3B and 3C). These findings indicate that IL-10 overexpression effectively prevented perivascular accumulation of inflammatory cells following SAH.

**Figure 3.**
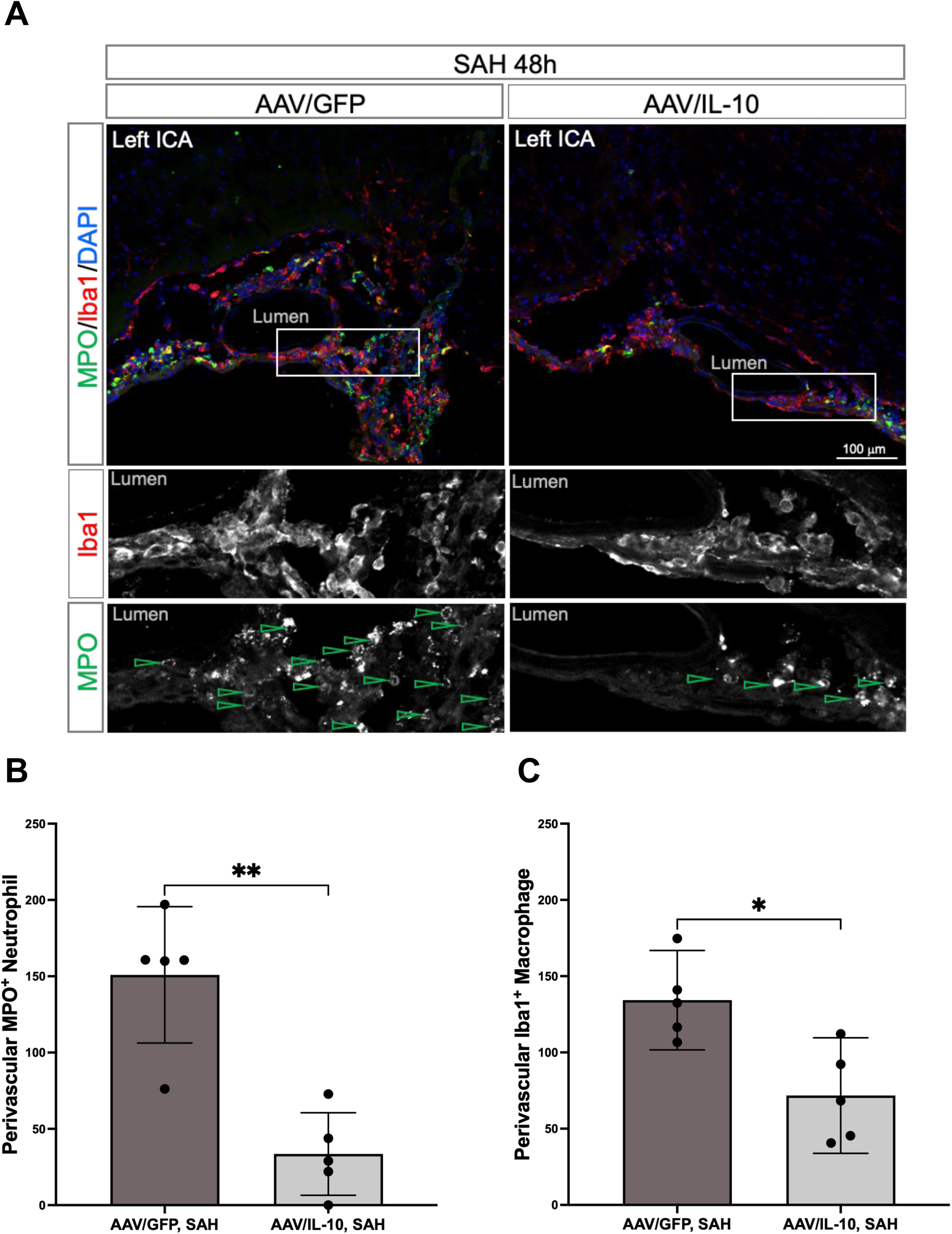
Interleukin-10 (IL-10) Reduces Perivascular Neutrophil and Macrophage Accumulation After Experimental Subarachnoid Hemorrhage (SAH). A, Immunohistochemical staining of sections containing the left internal carotid artery (ICA) obtained 48 h after SAH using the neutrophil marker myeloperoxidase (MPO) and the macrophage marker ionized calcium-binding adaptor molecule 1 (Iba1). The adeno-associated virus (AAV)/IL-10 SAH group exhibited reduced perivascular accumulation of neutrophils and macrophages compared with the AAV expressing green fluorescent protein (AAV/GFP) SAH group. B and C, Neutrophil and macrophage accumulation was significantly reduced in the AAV/IL- 10 SAH group compared with the AAV/GFP SAH group. Values are presented as mean ± SD, n= 5. \**P* < 0.05, \*\**P* < 0.01.

### IL-10 Suppressed SAH-Induced Increases in IL-6 and MMP-3

To identify key molecules involved in IL-10-mediated regulation of inflammatory pathways, a serum cytokine array was performed. The array performed 3 h after SAH showed significantly weaker MMP-3 signals in the AAV/IL-10 group than in the AAV/GFP group (Figure 4A). Using the same samples, ELISA confirmed that the AAV/IL-10 group exhibited significantly lower SAH-induced increases in MMP-3 (Figure 4B) and IL-6, a representative inflammatory cytokine (Figure 4C). These findings suggest that IL-10 overexpression suppresses the early systemic inflammatory response after SAH by downregulating key inflammatory mediators, including MMP-3 and IL-6.

**Figure 4.**
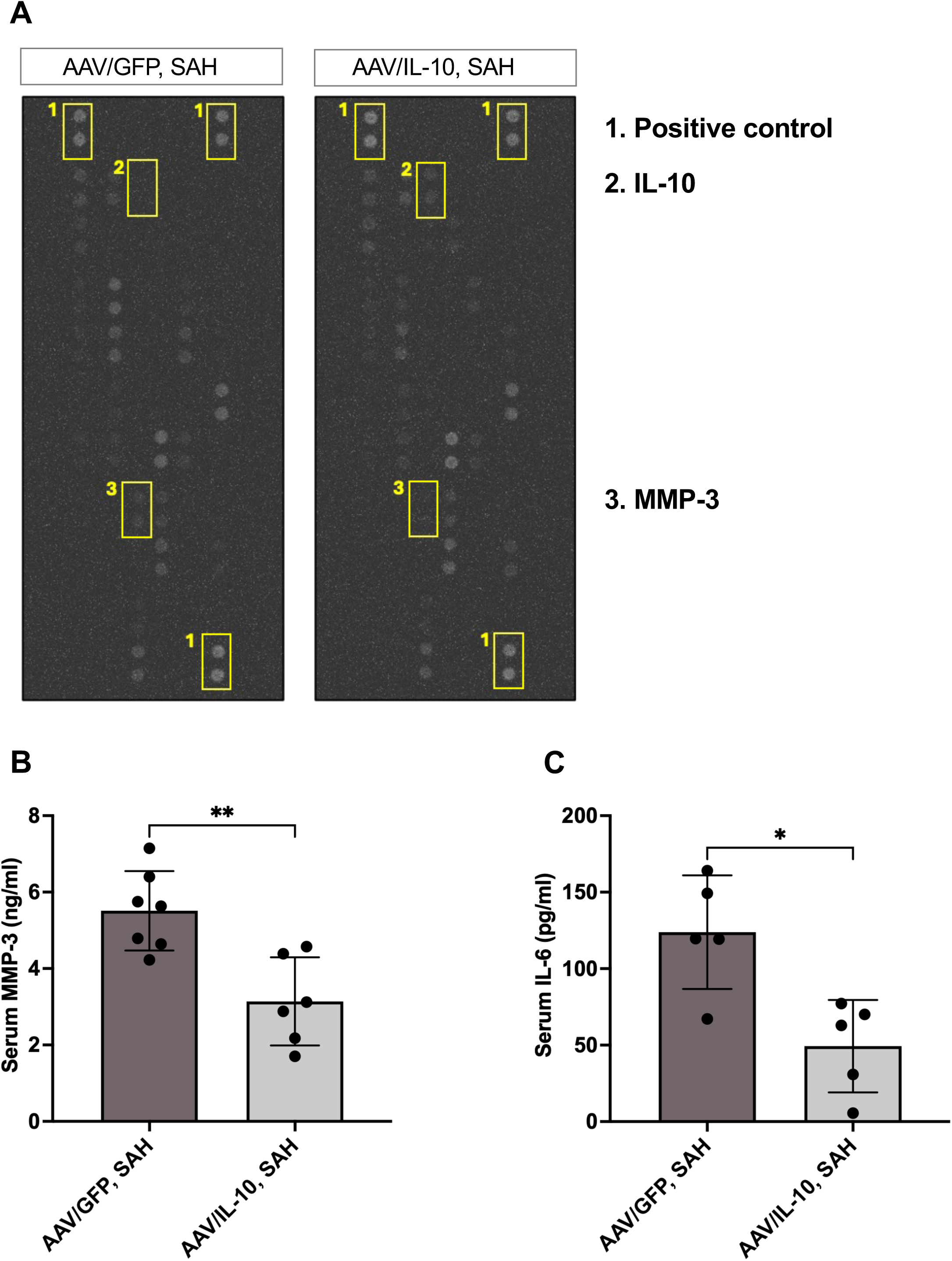
Interleukin-10 (IL-10) Suppresses Systemic Inflammatory Responses After Experimental Subarachnoid Hemorrhage (SAH). A, Representative cytokine array membranes showing serum levels of inflammatory and vascular cytokines in adeno-associated virus expressing green fluorescent protein (AAV/GFP) SAH and AAV/IL-10 SAH mice. Signals corresponding to IL-10 and matrix metalloproteinase- 3 (MMP-3) are shown. The intensity of the MMP-3 signal was reduced in the AAV/IL-10 SAH group. B and C, Serum concentrations of MMP-3 and interleukin-6 (IL-6) were measured by ELISA. Both cytokines were significantly reduced in the AAV/IL-10 SAH group compared with the AAV/GFP SAH group. Values are presented as mean ± SD, n = 5. \**P* < 0.05, \*\**P* < 0.01.

### AAV/IL-10 Vector Administration Remained Effective After SAH Induction

To evaluate the therapeutic potential of IL-10 overexpression in a clinically relevant setting, the AAV/IL-10 vector was administered immediately after SAH induction (Figure 5A). Neurological scores did not differ significantly from those in the AAV/GFP SAH group on day 1 but were significantly improved on days 2 and 3 after SAH (Figure 5B and 5C).

**Figure 5.**
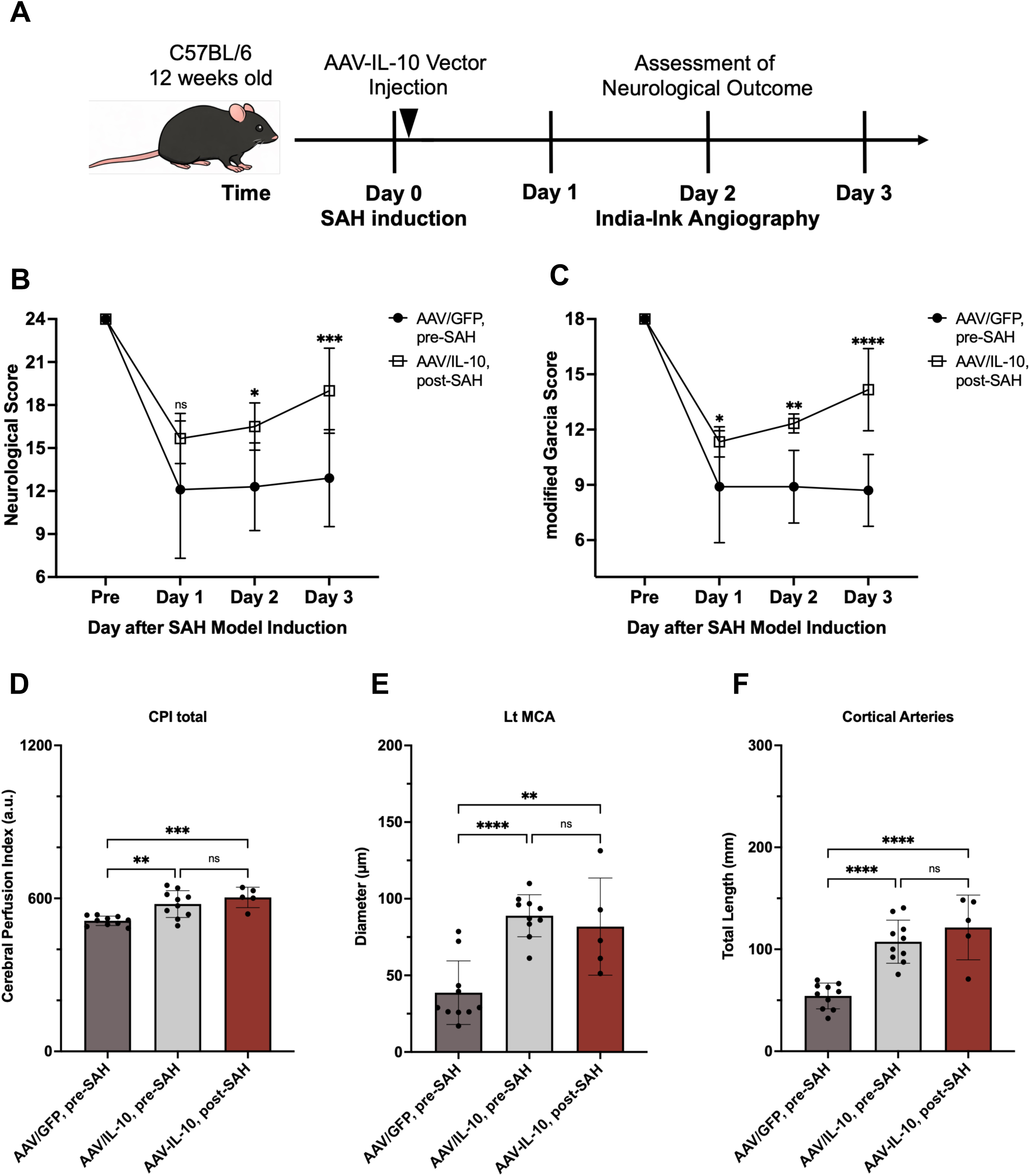
Post-Subarachnoid Hemorrhage (SAH) Administration of Adeno-Associated Virus Encoding Interleukin-10 (AAV/IL-10) Improves Neurological Outcomes, Cerebral Microcirculation, and Cerebral Vasospasm. A, Experimental protocol. The AAV/IL-10 vector was administered immediately after SAH induction. Neurological outcomes were evaluated for up to 3 days after SAH, and India ink angiography was performed 48 h after SAH. B and C, Neurological outcomes after SAH assessed using the neurological and modified Garcia scores. Neurological outcomes were significantly improved compared with those in the AAV expressing green fluorescent protein (AAV/GFP) SAH group. Values are presented as mean ± SD, n = 10 in the AAV/GFP SAH group and n = 5 in the AAV/IL-10 post-SAH group. \**P* < 0.05, \*\**P* < 0.01, \*\*\**P* < 0.001, \*\*\*\**P* < 0.0001. D–F, India ink angiography demonstrated that the cerebral perfusion index, left middle cerebral artery (Lt MCA) diameter, and total cortical arterial length were significantly improved in the AAV/IL-10 post-SAH group compared with the AAV/GFP SAH group, with no significant differences compared with the AAV/IL-10 SAH group. Values are presented as mean ± SD, n = 10 in the AAV/GFP SAH and AAV/IL-10 SAH groups and n = 5 in the AAV/IL-10 post-SAH group. \*\**P* < 0.01, \*\*\**P* < 0.001, \*\*\*\**P* < 0.0001.

India ink angiography performed 48 h after SAH showed that post-SAH administration of the AAV/IL-10 vector significantly improved the CPI, Lt MCA diameter, and total length of visible cortical arteries compared with the AAV/GFP SAH group (Figure 5D–F). These findings indicate that administration of the AAV/IL-10 vector after SAH onset attenuated cerebral vasospasm and microcirculatory impairment once systemic IL-10 expression reached an effective level. Overall, post-SAH administration of the AAV/IL-10 vector improved neurological outcomes, cerebral perfusion, MCA diameter, and cortical arterial filling compared with the AAV/GFP SAH group.

## DISCUSSION

The main finding of this study was that IL-10 overexpression improved neurological outcomes after SAH, highlighting its potential for clinical application. In a mouse model of SAH, IL-10 overexpression attenuated the decrease in cerebral perfusion and improved neurological function. These effects were accompanied by reduced accumulation of neutrophils and macrophages around the cerebral arteries and were associated with lower peripheral blood levels of MMP-3 and IL-6. These results suggest that IL-10 exerts neuroprotective effects following SAH through its anti-inflammatory properties. Furthermore, the AAV/IL-10 vector demonstrated efficacy even when administered after SAH onset. Consequently, AAV-mediated IL-10 delivery may represent a promising therapeutic strategy for SAH.

Several preclinical studies have reported that anti-inflammatory interventions improve neurological outcomes after SAH. For instance, INT-777, a Takeda G protein-coupled receptor 5 (TGR5) agonist, exerts neuroprotective effects by binding to TGR5 and suppressing inflammatory cytokine production and neutrophil migration following SAH.^34^ In addition, CDDO, a triterpenoid compound with anti-inflammatory properties,^35^ and FTY720, a sphingosine-1-phosphate receptor agonist,^36^ have shown neuroprotective effects by inhibiting inflammatory responses after SAH. However, a recent systematic review concluded that the current evidence is insufficient to establish the efficacy of anti-inflammatory interventions for SAH treatment.^37^ Therefore, we investigated sustained overexpression of the endogenous immunoregulatory cytokine IL-10 using an AAV vector. To our knowledge, this is the first study to demonstrate that IL-10 overexpression improves neurological outcomes after SAH through its anti-inflammatory effects, providing important evidence supporting the development of future anti-inflammatory therapies for SAH.

The beneficial effects of IL-10 overexpression on cerebral vasospasm and microcirculatory dysfunction may involve suppression of activated neutrophil and macrophage accumulation around the cerebral arteries after SAH. In the early phase after SAH, neutrophils migrate to the perivascular region, where they produce inflammatory cytokines and form NETs.^33,38–41^ IL-10 may inhibit neutrophil accumulation and the associated inflammatory response. Macrophages also migrate to the perivascular regions early after SAH, thereby amplifying inflammatory responses.^23,42^ The beneficial effects of IL-10 may be associated with attenuation of inflammation and reduced pro-inflammatory macrophage responses following neutrophil activation. Taken together, these findings suggest that the anti-inflammatory effects of IL-10 suppress these two immune cell populations, thereby attenuating cerebral vasospasm and microcirculatory dysfunction after SAH.

Following SAH, the production of various inflammatory cytokines is upregulated in multiple cell types, including immune and endothelial cells, forming a complex inflammatory cascade.^10,43^ In this study, serum concentrations of IL-6 and MMP-3 were lower in the AAV/IL- 10 SAH group. IL-6 is a representative inflammatory cytokine that activates immune cells, including neutrophils and macrophages, after SAH, thereby inducing further cytokine production and triggering a potent inflammatory response,^17,42^ which is strongly associated with neurological deterioration after SAH.^43–45^ MMP-3 is one of the MMPs induced by inflammatory cytokines and contributes to BBB disruption and increased vascular permeability after SAH.^46^ Moreover, MMP-3 levels are elevated in patients with cerebral vasospasm. Upregulation of these inflammatory mediators after SAH further amplifies inflammation and contributes to neuronal injury. Although the precise mechanisms remain to be fully elucidated, our findings suggest that IL-10 exerts neuroprotective effects by suppressing inflammatory cell migration and subsequent cytokine production, thereby preserving BBB integrity and ameliorating microcirculatory dysfunction.

This study also highlights the potential of the AAV/IL-10 vector as a novel therapeutic approach for SAH. Currently, drugs targeting the cerebral vasculature after SAH, such as Rho- kinase inhibitors and endothelin receptor antagonists, primarily target delayed cerebral vasospasm. However, these agents often fail to produce significant improvements in functional neurological outcomes.^47^ In contrast, the present study suggests that IL-10 overexpression ameliorates microcirculatory dysfunction driven by cerebrovascular inflammation, which begins much earlier than delayed cerebral vasospasm. By targeting this early phase, IL-10 may mitigate the sequential pathological cascade from EBI to DCI. Therefore, administration of the AAV/IL-10 vector may represent a promising therapeutic approach.

However, this study has some limitations. First, the detailed mechanisms by which IL-10 overexpression inhibits the SAH-induced inflammatory cascade remain unclear. It will be important to identify the signaling pathways through which IL-10 exerts its protective effects after SAH. Second, although IL-10 was overexpressed using a viral vector to reliably induce anti-inflammatory effects, sustained anti-inflammatory activity may alter metabolism, homeostasis, and immune regulation, potentially affecting clinical outcomes after SAH.^48^ Third, a time lag may exist before the AAV/IL-10 vector exerts its therapeutic effects because IL-10 expression requires time to reach effective systemic concentrations. Because early intervention against EBI is critical for neuroprotection after SAH, this delay may limit efficacy during the early phase. Although direct administration of recombinant IL-10 could provide immediate effects, its short half-life makes continuous suppression of the prolonged inflammatory cascade from EBI to DCI challenging. Therefore, a hybrid strategy combining acute administration of recombinant IL-10 protein to bridge the ultra-early temporal gap with AAV-mediated gene therapy for sustained neuroprotection may represent an optimal therapeutic approach. Further studies are warranted to elucidate the precise mechanisms of IL-10, determine the optimal concentration, and optimize the delivery strategy to achieve sufficient anti-inflammatory effects after SAH.

In conclusion, AAV-mediated IL-10 overexpression significantly improved neurological outcomes following experimental SAH. These neuroprotective effects were associated with attenuation of inflammatory responses, cerebral vasospasm, and microcirculatory dysfunction, as well as preservation of cerebral perfusion. These findings highlight the importance of early control of inflammation in determining neurological outcomes after SAH and support further investigation of IL-10-based therapeutic strategies.

## Acknowledgments

We gratefully acknowledge Tosoh Corporation for kindly providing the AVR gel. We extend our appreciation to the technicians involved in vector production, including Ken Sugo, Hiroto Kodera, Masumi Nishina, Yui Komaki, and Azusa Onodera. We also thank Anri Machi and Takashi Tamatani of Kanazawa University for their technical support

## Nonstandard Abbreviations and Acronyms

SAH: subarachnoid hemorrhage
ICA: internal carotid artery
MCA: middle cerebral artery
DCI: delayed cerebral ischemia
EBI: early brain injury
IL-10: interleukin-10
AAV: adeno-associated virus
GFP: green fluorescent protein
RAGE: receptor for advanced glycation end product
BBB: blood–brain barrier
MPO: myeloperoxidase
CPI: cerebral perfusion index
MMP-3: metalloproteinase-3
ANOVA: analysis of variance
SD: standard deviation.

## Major Resources Table

In order to allow validation and replication of experiments, all essential research materials listed in the Methods should be included in the Major Resources Table below. Authors are encouraged to use public repositories for protocols, data, code, and other materials and provide persistent identifiers and/or links to repositories when available. Authors may add or delete rows as needed.

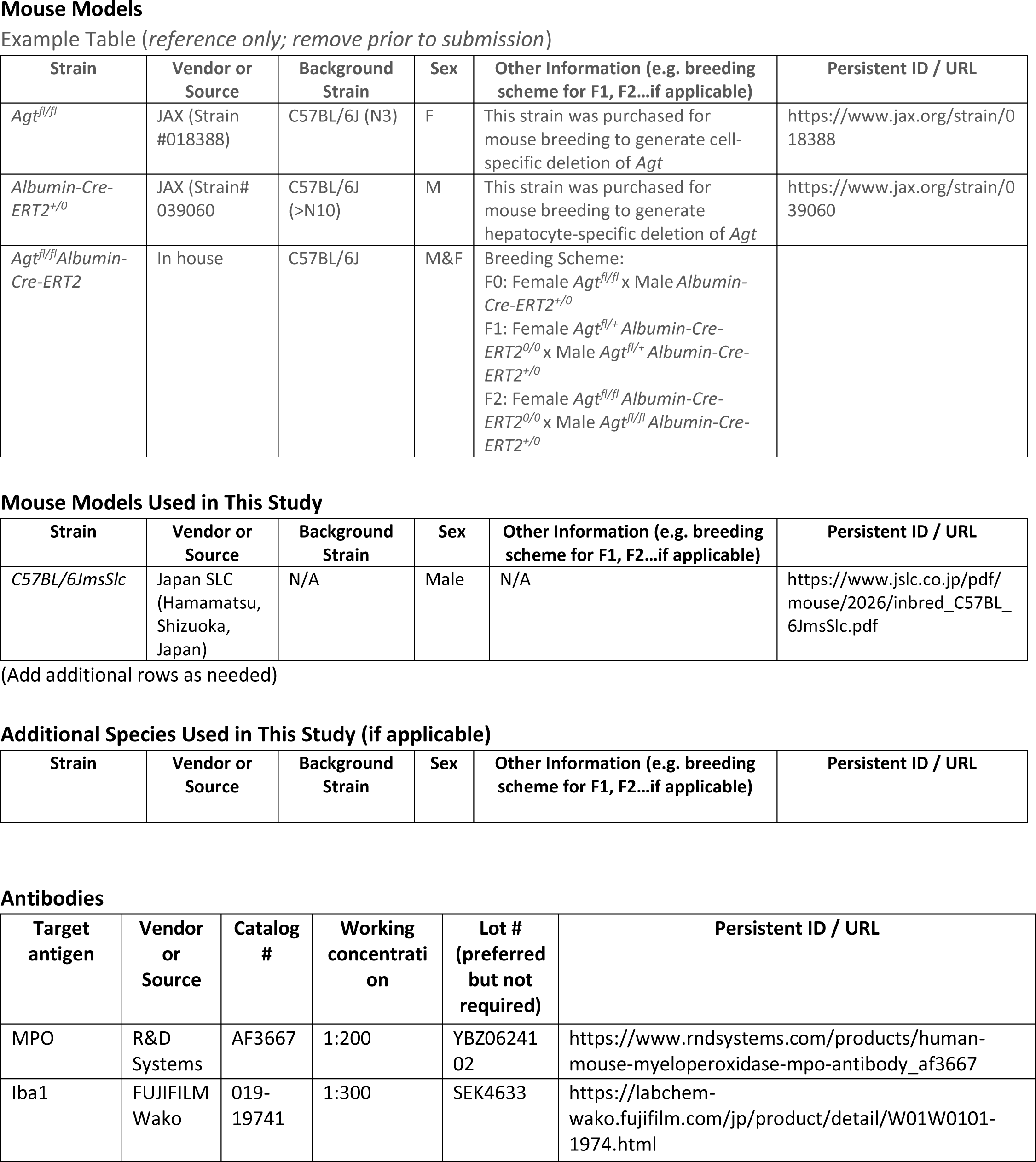
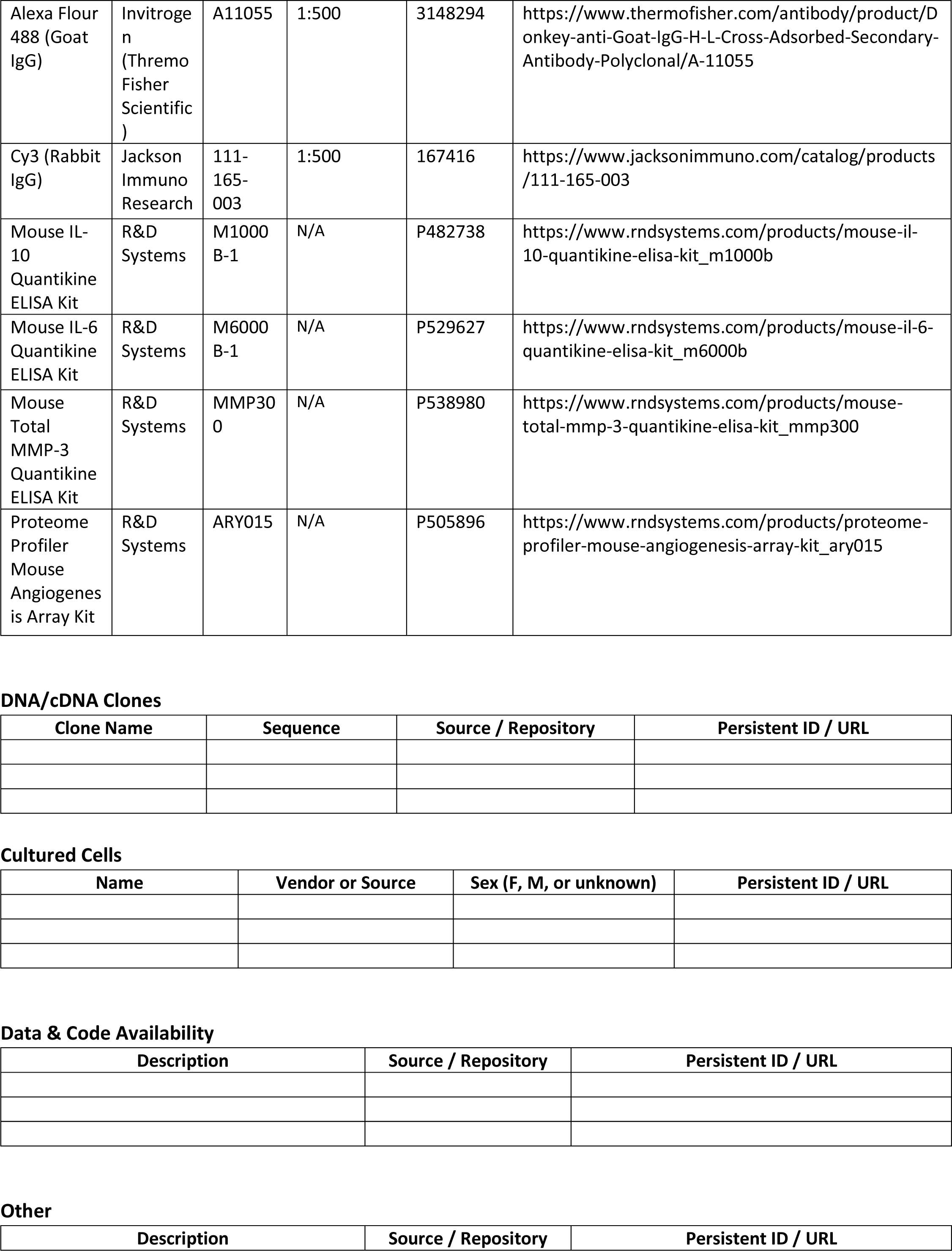
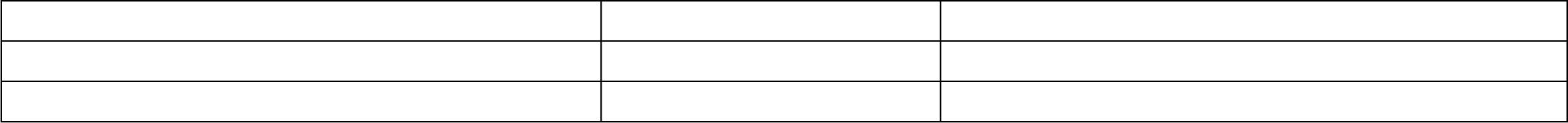

## ARRIVE GUIDELINES

The ARRIVE guidelines (https://arriveguidelines.org/) are a checklist of recommendations to improve the reporting of research involving animals. Key elements of the study design should be included below to better enable readers to scrutinize the research adequately, evaluate its methodological rigor, and reproduce the methods or findings.

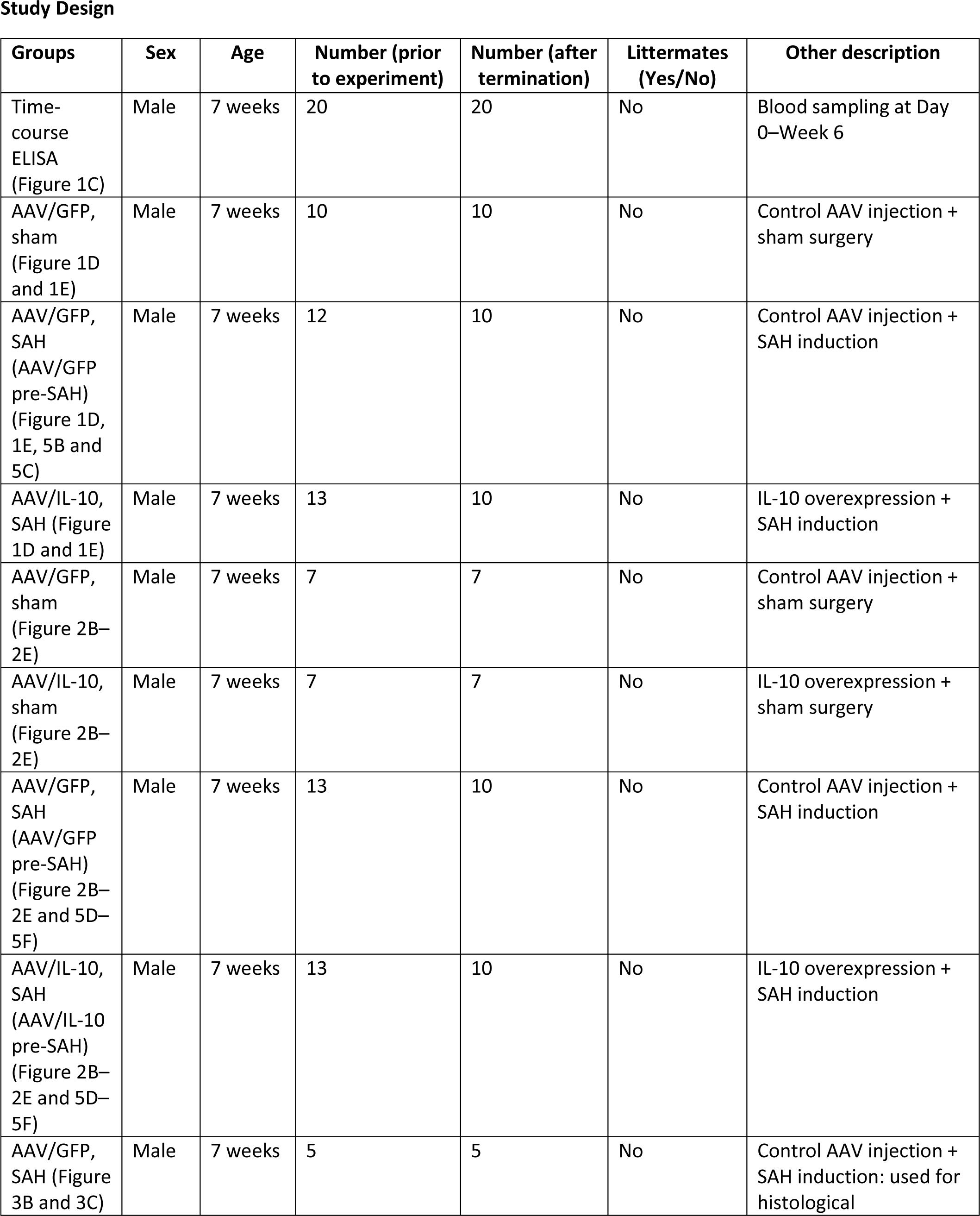
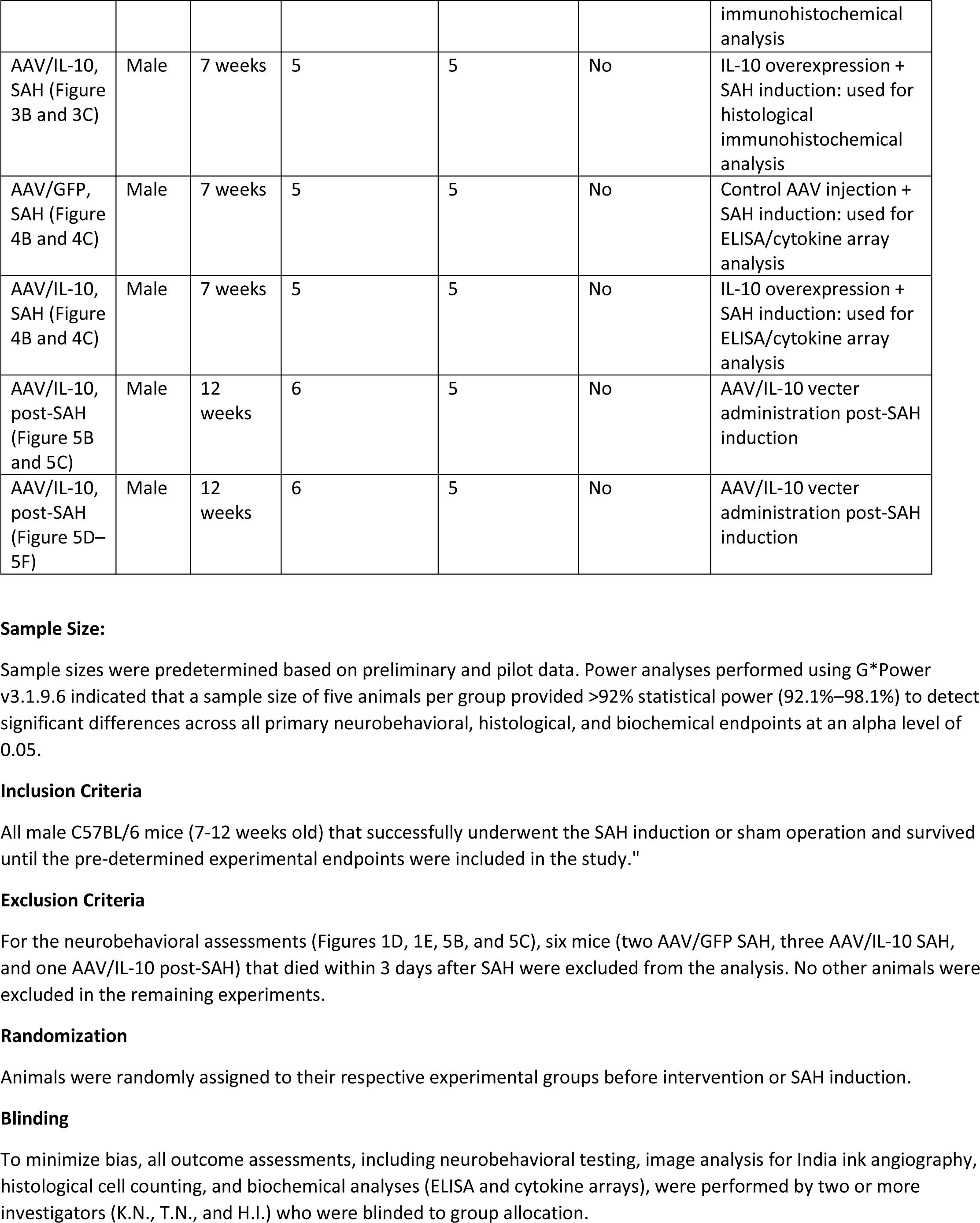

Study Design
| Groups | Sex | Age | Number (prior to experiment) | Number (after termination) | Littermates (Yes/No) | Other description |
| --- | --- | --- | --- | --- | --- | --- |
| Time-course ELISA (Figure 1C) | Male | 7 weeks | 20 | 20 | No | Blood sampling at Day 0–Week 6 |
| AAV/GFP, sham (Figure 1D and 1E) | Male | 7 weeks | 10 | 10 | No | Control AAV injection + sham surgery |
| AAV/GFP, SAH (AAV/GFP pre-SAH) (Figure 1D, 1E, 5B and 5C) | Male | 7 weeks | 12 | 10 | No | Control AAV injection + SAH induction |
| AAV/IL-10, SAH (Figure 1D and 1E) | Male | 7 weeks | 13 | 10 | No | IL-10 overexpression + SAH induction |
| AAV/GFP, sham (Figure 2B–2E) | Male | 7 weeks | 7 | 7 | No | Control AAV injection + sham surgery |
| AAV/IL-10, sham (Figure 2B–2E) | Male | 7 weeks | 7 | 7 | No | IL-10 overexpression + sham surgery |
| AAV/GFP, SAH (AAV/GFP pre-SAH) (Figure 2B–2E and 5D–5F) | Male | 7 weeks | 13 | 10 | No | Control AAV injection + SAH induction |
| AAV/IL-10, SAH (AAV/IL-10 pre-SAH) (Figure 2B–2E and 5D–5F) | Male | 7 weeks | 13 | 10 | No | IL-10 overexpression + SAH induction |
| AAV/GFP, SAH (Figure 3B and 3C) | Male | 7 weeks | 5 | 5 | No | Control AAV injection + SAH induction: used for histological immunohistochemical analysis |
| AAV/IL-10, SAH (Figure 3B and 3C) | Male | 7 weeks | 5 | 5 | No | IL-10 overexpression + SAH induction: used for histological immunohistochemical analysis |
| AAV/GFP, SAH (Figure 4B and 4C) | Male | 7 weeks | 5 | 5 | No | Control AAV injection + SAH induction: used for ELISA/cytokine array analysis |
| AAV/IL-10, SAH (Figure 4B and 4C) | Male | 7 weeks | 5 | 5 | No | IL-10 overexpression + SAH induction: used for ELISA/cytokine array analysis |
| AAV/IL-10, post-SAH (Figure 5B and 5C) | Male | 12 weeks | 6 | 5 | No | AAV/IL-10 vector administration post-SAH induction |
| AAV/IL-10, post-SAH (Figure 5D–5F) | Male | 12 weeks | 6 | 5 | No | AAV/IL-10 vector administration post-SAH induction |

